# Preserved synaptic architecture but impaired ketamine-induced synaptic plasticity of layer 5 pyramidal neurons in the aged frontal cortex

**DOI:** 10.1101/2025.09.02.673642

**Authors:** IF Ugidos, JC Iglesias, S Milanes, F Farahani, R. Mostany

**Affiliations:** Pharmacology Department, Tulane School of Medicine. Tulane University, New Orleans, LA (USA); Tulane Brain Institute. Tulane University, New Orleans, LA (USA); Tulane Medical Program. Tulane University, New Orleans, LA (USA)

**Keywords:** Aging, dendritic spines, *in vivo* imaging, frontal cortex, ketamine

## Abstract

Healthy aging is accompanied by a gradual decline in higher-order cognitive functions, including working memory, attention, and cognitive flexibility, processes that critically rely on intact frontal cortical circuits. While neuronal loss is minimal during aging, whether there are changes in functional plasticity in this region remains unexplored. In this regard, dendritic spines, the primary postsynaptic structures of excitatory synapses, act as key hubs for experience-dependent synaptic remodeling. Using longitudinal *in vivo* two-photon imaging in Thy1-eGFP-M mice, we examined age-related changes in dendritic spine density and dynamics in layer 5 pyramidal neurons of the secondary motor area (MOs), a frontal cortical region essential for strategy switching and cognitive flexibility, and that was assessed using an operant conditioning paradigm.

We found that aged mice (18–22 months) exhibited significant impairments in cognitive flexibility relative to young mice (3–5 months) in the four-odor choice discrimination and reversal task. Analysis of dendritic spine plasticity revealed that baseline spine density, turnover, and morphology were largely preserved in aged mice. Sex differences were evident, with females displaying higher spine density and a greater fraction of stable spines, a feature maintained across aging. Importantly, despite preserved baseline architecture, aged mice showed impaired ketamine-induced spinogenesis and reduced stabilization of newly formed spines, in contrast to the robust structural plasticity observed in young mice.

These results indicate that healthy aging selectively impairs activity-dependent synaptic remodeling without affecting steady-state spine architecture in frontal cortical circuits. By linking deficits in induced synaptic plasticity to age-related impairments in cognitive flexibility, our study highlights the critical need to target plasticity mechanisms as a therapeutic strategy to restore executive function and cognitive adaptability in the aging brain.

## Introduction

Healthy aging is accompanied by a decline in several higher-order cognitive domains, including working memory, attention, and cognitive flexibility^1,2^, that are tightly associated with the correct functioning of the frontal cortical region^3–5^. Although it was believed that those deficits could arise from neuronal loss, it has been demonstrated that neurons are fairly preserved during aging^6,7^. Current studies suggest that, rather than neuronal loss, the aging brain presents decreased functional plasticity—the dynamic modulation of synaptic connections in response to experience—correlating with the onset of age-related cognitive deficits^8–11^.

Dendritic spines, the primary postsynaptic structures of excitatory synapses in the mammalian brain, represent a central hub for synaptic plasticity^12^. While some dendritic spines remain stable over long periods of time, a significant fraction of them is continuously being formed and eliminated—a process that enables experience-dependent learning and flexible circuit reorganization^13^. These dynamic changes lead to input rearrangements that can have profound effects on network function, and thus highlighting the critical role of spine dynamics in shaping brain connectivity^14^. Traditional studies using fixed tissue have provided valuable information regarding the density of dendritic spines^15,16^, however, these studies lack the longitudinal examination of individual dendritic spines that would allow to capture the dynamic features of these synaptic structures. In this context, *in vivo* longitudinal imaging represents as a powerful tool to track spine dynamics over time and reveal ongoing synaptic remodeling in the living brain^17,18^.

In this study we investigated the impact of aging in the dynamic regulation of dendritic spine plasticity in the frontal cortex. We focus on the secondary motor area (MOs), a region critically involved in strategy shifting and decision making, key components of cognitive flexibility^19–23^ that have been demonstrated to be impaired with aging^24,25^. Previous studies using fixed tissue have reported conflicting results regarding whether aging leads to synaptic loss in the frontal cortex^10,15,26,27^. Here, we specifically examined age-related changes in the density and dynamics of layer 5 pyramidal neurons within the MOs. In addition, we assessed the capacity for induced plasticity in this region by the administration of a single, low, subanesthetic dose of ketamine, which has been shown to reliably promote spinogenesis in layer 5 pyramidal neurons of the MOs^28,29^. To our knowledge, this is the first study to examine dendritic spine density and dynamics within the MOs under both baseline conditions and following a plasticity-inducing intervention in the aged brain.

## Methods

### Animals

Both male and female Thy-1 eGFP-M mice (The Jackson Laboratory, 007788, Tg (Thy1-EGFP)MJrs/J^30^)aged 3-5 months (young) and 18-22 months (aged) were used for these experiments. Mice were group-housed under a 12-hour light/dark cycle and food and water were provided *ad libitum*. All procedures were approved by the Tulane University Institutional Care and Use Committee, and were performed in accordance with the NIH Office of Laboratory Animal Welfarés Public Health Service Policy on Humane Care and Use of Laboratory Animals and Guide for the Care and Use of Laboratory Animals.

### Four-odor choice discrimination and reversal task

Differences in behavioral flexibility between young and aged mice were assessed using a 4-odor choice discrimination and reversal task. The task was implemented as previously described^31,32^ with minor modifications. Mice were food-restricted for four days (young) or five days (aged) prior to training. Each day, animals received 0.75 g of food per gram of body weight, typically administered at 10 AM. Food allotments were adjusted daily to ensure weight loss per day did not exceed 10% of the baseline body weight. Mice were weighed daily, and food restriction was considered successful once animals reached 80–85% of their initial weight.

On the first day of pre-training, mice were habituated to the arena and pots. Chocolate-flavored pellets (45 mg/each; Bio-Serv) were used as a reward and were placed in empty pots across the four arena quadrants. The mouse was placed in the central start cylinder, then allowed to explore and consume the pellets for 10 minutes. This was repeated three times for a total duration of 30 minutes.

On the second day of pre-training, mice were trained to dig for buried rewards. Trials progressed in difficulty: the first two trials were done without shavings, then two trials with light dusting followed by two trials with pots ¼ full, two ½ full, and four with the bait fully buried in shavings. Pot location was randomly alternated each trial. Trials were untimed. However, mice taking over 10 minutes to dig and eat the pellets were retrained the next day. Mice failing to learn after retraining were excluded.

The discrimination phase was carried out on day 3. Wood shavings were scented on the day of testing with anise (undiluted; McCormick), clove (1:10 in mineral oil; Artizen), litsea (1:10; Eden Gardens), and thyme (1:5; Plant Therapy) extracts at a concentration of 0.01 mL/g and dispensed with a spray bottle. Each pot contained one of the scented shavings, with anise paired with the food reward. Odor placement was pseudo-randomized to avoid repetition in the same quadrant across trials. In each trial, once the starting cylinder was lifted, the animal was allowed to explore the arena. A choice was defined as active digging in a pot (not chewing or smelling the shavings). After a choice was made, the cylinder was lowered to prevent further exploration. In the case of an incorrect choice, the trial was terminated and the mouse was gently returned to the start cylinder. If the choice was correct, the mouse was allowed to consume the reward, and the animal was gently returned to the starting cylinder. Pots were removed and re-baited after each trial. Trials with no choice within 3 minutes were recorded as omissions. Two consecutive omissions required a retraining trial with a baited pot of unscented shavings placed in the start cylinder. Omissions were excluded from the final analysis. Mice reached criterion upon completing 8 correct responses out of 10 consecutive trials. Once criterion was met, animals were returned to their home cage and provided with additional food to complete their daily assigned portion.

On the following day, mice first completed a recall of the initial odor discrimination to criterion in order to assure the memory was retained. Immediately after reaching criterion, the reversal task was initiated. For this phase of the task, baited odor was switched from anise to clove. A novel odor cue was also introduced, switching thyme with eucalyptus (1:10 in mineral oil, 0.01 mL/g of wood shavings; Plant Therapy). Mice were required to complete the task as described for discrimination until criterion was reached. Choices were recorded and categorized as follows: anise: perseverative error; eucalyptus: novel error; litsea: irrelevant error; clove: correct. They were categorized into exploitative choices (perseverative errors, returning to the previously rewarded odor) and exploratory choices (novel and irrelevant error, as well as correct choices, adaptive behavior of exploring novel odors in pursuit of a reward). Exploit index (the proportion of exploit choices^32^) was calculated as (# of exploit choices-# of explore choices)/total trials.

### Cranial window surgery procedure

The implantation of glass-covered cranial windows was performed as previously described^33^, with minor modifications to adapt the procedure to the MOs region^29,32^. A subset of mice received two cranial windows—one over the MOs region and one over the primary somatosensory cortex barrel field (S1BF)—to investigate the potential differential effects of ketamine on dendritic spine dynamics in these two distinct brain areas. Briefly, animals were anesthetized with isoflurane (5.0% for induction, 1.5% for maintenance in 0.6 L/min of O₂). To reduce inflammation and prevent swelling, mice were subcutaneously injected with carprofen (5.0 mg/kg bw) and dexamethasone (0.2 mg/kg body weight) before any incision was made. A 3 mm diameter circular craniotomy was made directly above the MOs region (centered at AP: +2, ML: 0) using a pneumatic drill. After bone removal and hemostasis using Gelfoam (Pfizer) soaked in sterile saline, a 3 mm diameter glass coverslip (#1; Electron Microscopy Sciences) was implanted over the intact dura and secured with ethyl-cyanoacrylate glue. For mice undergoing double cranial window implantation, once the MOs window was secured, a second 4 mm diameter craniotomy was performed over the somatosensory cortex (centered at AP: –1.9, ML: +3), and a 5 mm glass coverslip was positioned over the skull and affixed with ethyl-cyanoacrylate glue. The remaining exposed skull was then covered with dental acrylic (Lang Dental Mfg. Co., Inc.). A custom-made titanium head bar (9.5 × 3.2 × 1.1 mm) was embedded in the acrylic to allow head fixation during *in vivo* imaging sessions. Mice were allowed to recover for two to three weeks before the first imaging session.

### In vivo two-photon laser scanning microscopy

Two-photon laser scanner microscopy (2PLSM) imaging was performed with a custom-built microscope equipped with a Ti:Sapphire laser (Chameleon Ultra II; Coherent Inc.) tuned to 910 nm, and a 40X 0.8 NA water immersion objective (Olympus). Animals were anesthetized with isoflurane (5.0% for induction, 1.0–1.5% for maintenance) throughout each imaging session (20-50 min). Collection of images was performed using ScanImage 3.8 software. Layer 5 pyramidal neurons in the MOs region were identified within the coordinates of +1 to +3 mm anterior and ± 1 mm lateral to bregma, and confirmed by the presence of a cell body located at a depth of at least 400 μm. For the analysis of the density and dynamics of dendritic spines, 4 to 8 dendritic branches were imaged at high magnification (512 × 512 pixels, 0.152 µm/pixel, 1.5 µm z-steps). One to two neurons were imaged per mouse. The same dendritic fragments were located using a coordinate system and imaged across days (day 0, 4, 8, 12, and 16 for the spine dynamics study). All imaging sessions were performed following the same hour schedule (from 9 am to 2 pm), to avoid potential differences associated with the circadian cycle.

### Ketamine treatment

Mice were randomly assigned either to the control or to the ketamine-treated groups. Treated mice received a subanesthetic dose of ketamine hydrochloride (10 mg/kg bw, i.p., Zetamine, VetOne) previously diluted in sterile saline to 1 mg/mL concentration on day 8, immediately after the third baseline *in vivo* imaging session. Control mice received the equivalent volume of sterile saline.

### Analysis of dendritic spine density and dynamics

Dendritic spine density and dynamics were analyzed using a custom MATLAB-based spine analysis software. All visible spines were manually annotated based on the following criteria: lateral spines were required to extend from the dendritic shaft by a distance equal or greater than one-third of the dendrite shaft’s width, and spines projecting along the z-axis had to be clearly visible in at least two consecutive optical slices. A minimum number of 100 spines per neuron on day 0 was required for the data from each neuron to be included in the experiment. Baseline dendritic spine density and dynamics (gained, lost, turnover rate) were quantified between imaging sessions at timepoints 0, 4, and 8 days, i.e., timepoints before ketamine or saline treatment. Gained spines were defined as those that newly appeared since the previous imaging session, while lost spines were defined as those that disappeared between imaging sessions. Spine dynamics were quantified either as the number of gained and lost spines per 100 μm, or as the gained and lost fraction (number of gained or lost spines divided by the total number of spines). Turnover rate (TOR) is defined as the combined number of gained and lost spines per 100 μm from the previous imaging session ([#gained spines + #lost spines]/100 μm). Transient spines were defined as those that appeared and disappeared in consecutive imaging sessions, whereas new persistent spines (NPS) were those that were first detected on a given imaging time point, and were still present at the following imaging session. Predicted survival functions of dendritic spines were calculated by fitting the survival fractions at the time points of interest to a one-phase exponential decay, and their plateaus and the rate constants were compared between experimental groups. The 8-day survival fraction was calculated taking into account 3 timepoints, either pre-ketamine (d0, d4, and d8), or post-ketamine (d8, d12, d16). To analyze the effect of ketamine, densities and dynamics were compared between the baseline timepoints and post-ketamine or post-saline administration (days 12 and 16).

### Analysis of dendritic spine morphology

Semiautomated classification of dendritic spine subtypes was performed using ImageJ and a custom MATLAB routine, as previously described^18,34,35^. Given the limited z-resolution of 2PLSM images, the spine morphology analysis was restricted to laterally protruding spines whose maximum brightness occurred on the same optical slice as the one where the dendritic shaft is. For morphological evaluation, a 5-pixel-wide line was drawn along each spine, extending from its base at the dendritic shaft to the tip of the spine. The underlying brightness intensity profile was extracted, and the MATLAB routine calculated the first derivative function (Df) of each pixel intensity trace. Morphological classification was then automatically assigned based on the number of instances that Df(x) crossed y = 0: derivative functions for a trace with no zero-crossings were classified as stubby, those with one crossing as mushroom, and those with two or more crossings as thin. For instance, the Df(x) of the brightness intensity profile of a mushroom dendritic spine typically crosses y = 0 once at the neck and again near the center of the spine head. A total of 50–60 spines per neuron were analyzed.

### Apical dendritic tree reconstruction and analysis

Reconstruction of apical dendritic trees of layer 5 pyramidal neurons was performed in 2PE z-stacks (100 optical slices, 5 µm z-step, 512×512 pixels, 0.72×0.75 μm/pixel) by using the SNT’s Reconstruction Viewer plugin for ImageJ^36^. Only neurons that displayed a complete apical dendritic tree with no shadows from brain vasculature were utilized for this analysis. Briefly, every dendritic tree was scaled, and semiautomatically traced from the deepest slice to the most superficial one. The number and length of dendritic branches and their boundary size was quantified.

### Statistical analyses

Analysis was performed using GraphPad Prism (GraphPad Software, LLC). Analysis of the results from the behavior studies was performed using a two-way ANOVA (# trials to criterion, type of error, strategy) or *t*-test (exploit index and # trials to first exploration). Statistical analyses of baseline dendritic spine density and dynamics between young and aged mice were done using *t*-test, while sex-differences between young and aged mice were performed using two-way ANOVA followed by a multiple comparison analysis (Fisher’s LSD when no interaction between variables, and Tukey’s *post hoc* correction when sex and age variables interact). The effect of ketamine on density, gain, loss, and 8-day survival fraction of dendritic spines between experimental groups was compared using two-way ANOVA, or a paired two-way ANOVA with repeated measurements for the comparison of baseline vs treatment for the analysis of transient and new persistent spines. Survival functions were statistically compared using the extra-sum-of-squares *F* test taking into account the plateau as well as the decay constant (*K*). Significance was set at *p* < 0.05.

## Results

### Cognitive flexibility is impaired in aged mice

We assessed age-related changes in cognitive flexibility using a 4-choice odor discrimination and reversal task (Fig. 1A and B). We selected this task because it is known to depend on the MOs during the reversal phase^31,32^, allowing us to target the same region accessible through a cranial window for studying *in vivo* structural plasticity dynamics^32^. We found that, while young and aged mice needed similar number of trials to reach criterion during the discrimination phase, aged mice performed significantly worse during the reversal (Fig. 1C), requiring more trials to meet criterion and making significantly more perseverative errors (Fig. 1C and D). As a consequence, aged mice displayed a higher number of exploitation trials relative to exploration trials, as well as compared to exploitation trials in young mice (Fig. 1E), resulting in an exploitation index that tends to be higher in aged compared to young mice (Fig. 1F). The impaired behavior flexibility is also evidenced by the significant higher number of trials needed by the aged mice to switch from an exploitative (based on previously rewarded odor) to an exploratory (based on exploration of new odors to find the reward) foraging strategy (Fig. 1G). These results indicate a decreased cognitive flexibility in aged mice compared to young mice.

**Figure 1.**
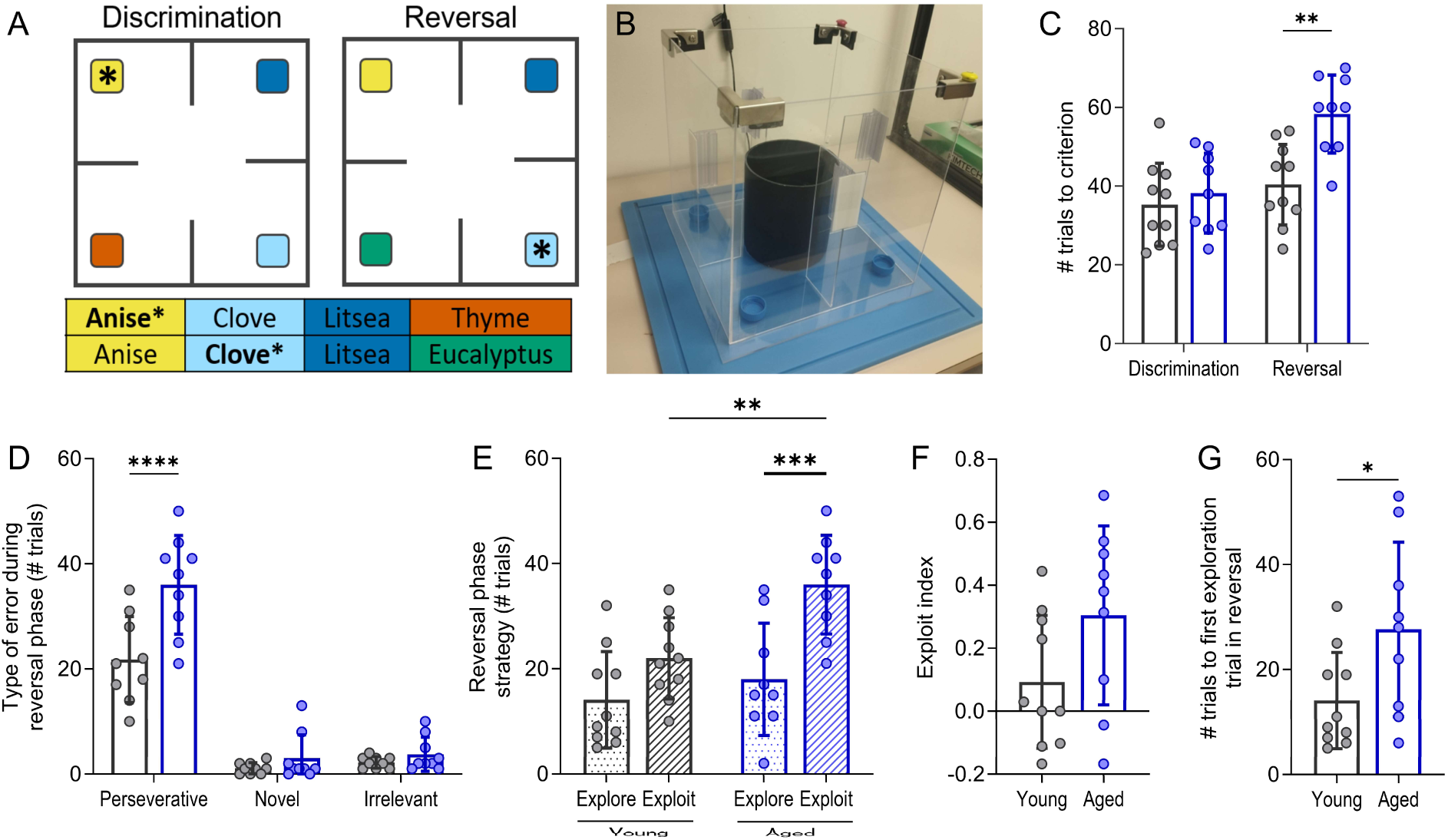
**A** Representation of the arena with the odor choices for the discrimination and reversal phases of the task. Asterisk shows baited odor. **B** Picture of the behavioral arena. **C** Deficits in behavior flexibility in aged compared to young mice are detected during the reversal phase (Two-way ANOVA: Age X Phase F(1,34) = 5.097, *p* = 0.035, Tukey’s post-hoc test *p* = 0.029). **D** Aged mice showed an increased number of perseverative errors compared to young mice (Two-way ANOVA: Age X Choice F(2, 48) = 7.516, *p* = 0.0014, Tukey’s post-hoc test: *p* < 0.0001). **E** This increase in perseverative errors in aged mice represents an increase of the exploitation versus exploration strategy as well as an increase in exploitation in aged compared to young mice (Two-way ANOVA Age X Strategy F(1,34) = 2.822, *p* = 0.1021, Fisher LSD: exploitation young vs exploitation aged: *p* = 0.0023; exploration vs exploitation in aged: *p* = 0.0002). **F** This shift in strategy with aging is shown by a trend to an increased exploit index (Unpaired t-test: *p* = 0.0812), **G** and it is supported by the increase in the number of trials needed by aged mice to run the first exploration trial in the reversal phase (Unpaired t-test: 0.0392). n = 10 young and n = 9 aged mice. Data presented as mean ± SD.

### Steady-state spine dynamics are preserved in aged mice

Following our observations of diminished cognitive flexibility with aging, we sought to determine whether synaptic plasticity deficits were present in the MOs, a brain area involved in this task^31^, of aged mice. For this purpose, we analyzed the impact of aging on baseline, steady-state, spine dynamics of the apical tuft of layer 5 pyramidal neurons in the MOs region (Fig. 2A), since excitatory neurons in this cortical area are crucial for adaptative strategy learning in response to a rule shift^19,20,37^. We found no significant differences between young and aged mice in baseline dendritic spine density (Fig. 2B) or dynamics (Fig. 2C). The calculated survival fraction based on day 0 to day 16 imaging timepoints did not differ between age groups (Fig. 2D), neither did the proportions of the different type of dendritic spines (thin/stubby/mushroom; Fig. 2E). We also reconstructed the apical dendritic trees of these layer 5 pyramidal neurons (Fig. 2F), and we found that although the number of dendritic branching points was not significantly different between young and aged mice (Fig. 2G), the average length of the branches was reduced in aged compared to young mice (Fig. 2H), and most probably driving the trend to a reduced boundary size of the apical tuft (Fig. 2I). Altogether, these data indicate that the steady-state spine density, dynamics, and morphology are preserved in this cell type and brain region in aged mice.

**Figure 2.**
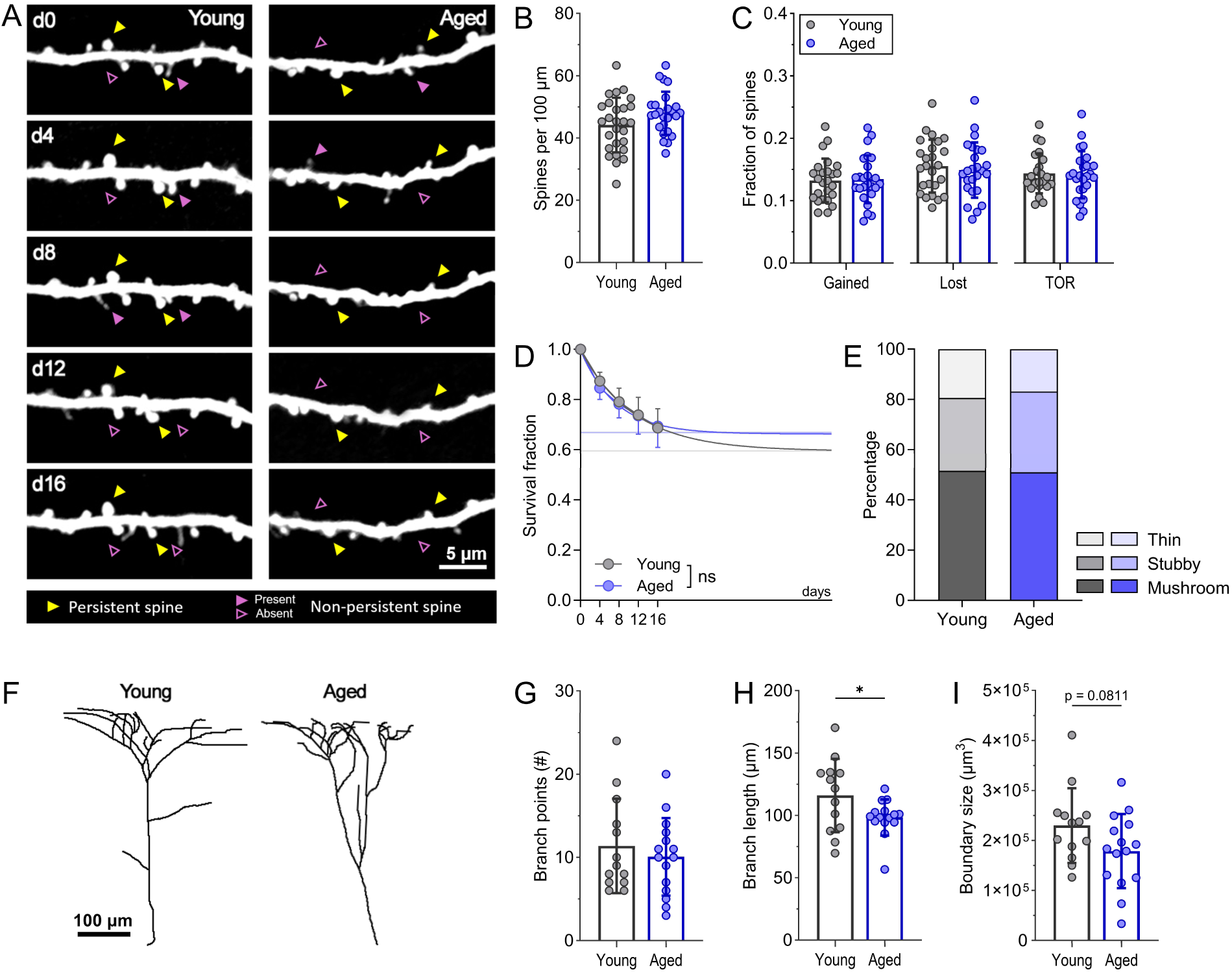
**A** Representative image of dendritic fragments of layer 5 pyramidal neurons in the MOs of young and aged mice across timepoints. **B** The density of dendritic spines is not altered by aging **C** nor their dynamics (gained, lost, turnover ratio). **D** The estimated fraction of persistent spines did not differ between young and age mice, **E** neither the percentages of the different spine morphologies (thin, stubby, mushroom). **F** Representative images of apical dendritic arbors reconstruction. **G** The number of dendritic branches of layer 5 pyramidal neurons did not differ between young and aged mice, **H** but the average branch length was significantly decreased in aged compared to young mice (Unpaired *t*-test: *p* = 0.050). **I** The boundary size of the apical arbor of the layer 5 pyramidal neurons analyzed was not significantly different between young and aged mice, although a trend was observed (Unpaired *t*-test: *p* = 0.0811), probably driven by the decrease of average branch length. Density, dynamics, and morphology: n = 31 neurons from 19 young mice, and n = 25 neurons from 15 aged mice. Branch analysis: n = 13 neurons young and n = 15 from aged mice. Survival function analysis: n = 12 neurons from 10 young mice and n = 10 neurons from 7 aged mice. Data presented as mean ± SD.

### Sex differences in steady-state spine density and dynamics are largely age-independent

Sex differences in dendritic spine density have been reported in mice^38^ and human^27^ cortices, therefore we aimed to investigate if sex differences were present in the MOs region and whether they were age-dependent (Fig. 3). We found that baseline spine density was significantly lower in male compared to female mice in both young and aged groups (Fig. 3B). Interestingly, aging led to an increase in spine density in males but not in females. Spine dynamics (gained, lost, TOR) were significantly higher in males compared to females when analyzing the fraction of spines (Fig. 3C). Interestingly, the dynamics observed in absolute number of spines per 100 μm show no difference within sexes (Suppl. Fig. 1), suggesting that dendritic spines of females are more stable than those of male mice. To confirm this finding, we analyzed the fraction of spines present at day 0 that persisted through day 16, and we found that females show higher stable fractions than male mice (Fig. 3D). We also fit the dendritic spine survival fractions from day 0 to days 4, 8, 12, and 16 to an exponential decay and found that females show a higher predicted survival fraction than males in both young and aged groups. Although the difference remained significant, it was reduced with aging (Fig. 3E). Consistent with our pooled data (Fig. 2D), no differences in predicted survival fractions were found between age groups. Dendritic spine morphology remains unchanged between sexes, although some trends may indicate a potential re-distribution of the type of spine in females versus males, independent of the age (Fig. 2E, Suppl. Fig. 1).

**Figure 3.**
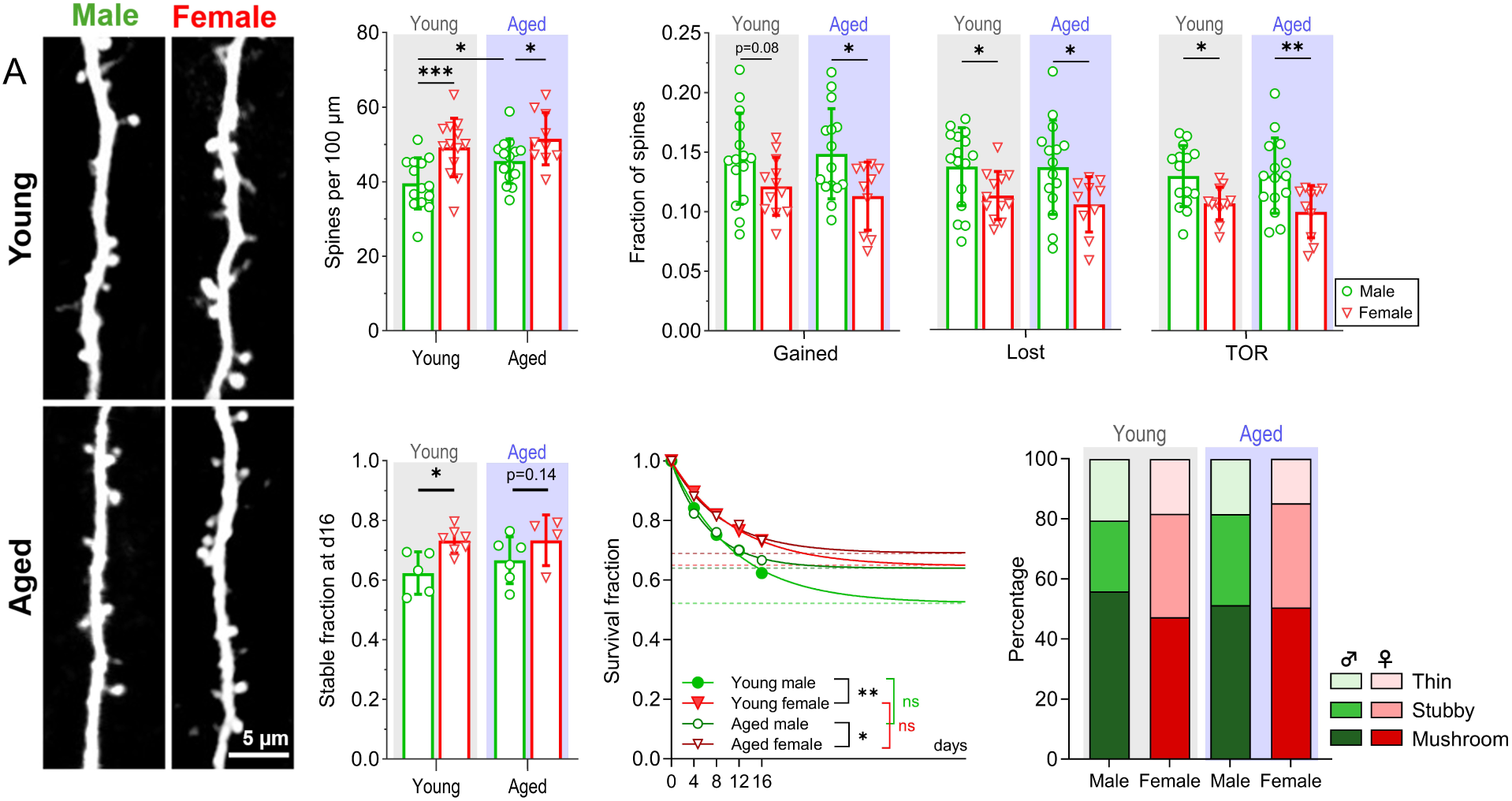
**A** Representative images of dendritic fragments of female and male, young and aged mice at day 0 of the imaging timeline. **B** Dendritic spine density of layer 5 pyramidal neurons in Mos is higher in females than in males (Two-way ANOVA: Sex F(1,48) = 16.44; *p* = 0.0002; Age F(1,48) = 4.62; *p* = 0.0367; Sex X Age F(1,48) = 0.88; *p* = 0.3531), displaying main factor sex differences in young (Fisheŕs LSD *p* = 0.0007) and aged (Fisheŕs LSD: *p* = 0.0372) mice. Aged males present higher spine density than young males (Fisheŕs LSD: *p* = 0.024), which is not happening in females. Sex differences in the metrics studied were not affected by age: fraction of gained spines (Two-way ANOVA: Sex F(1,48) = 9.625 *p* = 0.0032; Age F(1,48)=0.045 *p* = 0.83), fraction of lost spines (Two-way ANOVA: Sex F(1,48) = 9.159 *p =* 0.004; Age F(1,48)=0.41 *p* = 0.52), and TOR (Two-way ANOVA: Sex F(1,48) = 14.46 *p =* 0.0004; Age F(1,48)=0.21 *p* = 0.64). **D** The fraction of stable spines at day 16 (those that survived from day 0 to day 16) shows that females display higher stable fraction than males independently of the age (Two-way ANOVA: Sex F(1,18) = 8.76 *p =* 0.0084; Age F(1,18)=0.56 *p* = 0.46). **E** The estimated survival function indicates that the fraction of persistent dendritic spines is significantly different between males and females in both young (F test: *p* = 0.0002, plateau 0.52 in males vs 0.64 in females; *K* = 0.09 in females vs 0.09 in males), and aged (F test: *p* = 0.0069, plateau 0.69 in females vs 0.64 in males; *K* = 0.11 in females vs 0.15 in males). No aged-related differences were detected between young and aged mice within sexes. **F** Morphology of dendritic spines is mostly unaltered by sex or aging within sexes. Density and dynamics: n = 15/13 neurons from 11/10 male/female mice, and n = 15/11 neurons from 8/8 male/female aged mice. Survival fraction analysis: 5/7 neurons from 4/5 young mice and 6/4 neurons from 5/4 aged mice (males/females respectively in both young and aged mice). Data presented as mean ± SD.

### Ketamine-induced synaptic plasticity in the MOs region is hindered in the aged brain

Given the observed deficits in cognitive flexibility with aging despite preserved steady-state synaptic dynamics, and the fact that this behavioral task relies on synaptic plasticity in the MOs^32^, we hypothesized that aging may impair specifically activity-induced plasticity in this region. To elicit plasticity, we used ketamine, which has been shown to induce synaptic plasticity in pyramidal neurons in the MOs of young mice^29^ (Fig. 4A). Ketamine administration increased spine density in young mice by enhancing spine formation without altering spine elimination dynamics. Aged mice, however, did not show ketamine-induced changes in density or dynamics (Fig. 4B-D, Suppl. Fig. 2 A-E). Moreover, our data showed that both young female and male mice respond similarly to ketamine, exhibiting a comparable increase in spine density and in number of gained spines (Suppl. Fig. 2 F-H). When we analyzed the fate of the newly formed spines after ketamine administration, we found that the treatment increased both the number of transient spines as well as the number of new persistent spines (NPS) in young mice, but not in the aged group (Fig. 4 E,G). We found that the significant ketamine-induced increase in the number of transient spines in young males was not observed in female mice (Fig. 4F), which appears to be driven by a trend toward a higher baseline number of transient spines in young females compared to males. However, the primary effect of ketamine—the enhancement of newly persistent spines—is observed in both sexes (Fig. 4H).

**Figure 4.**
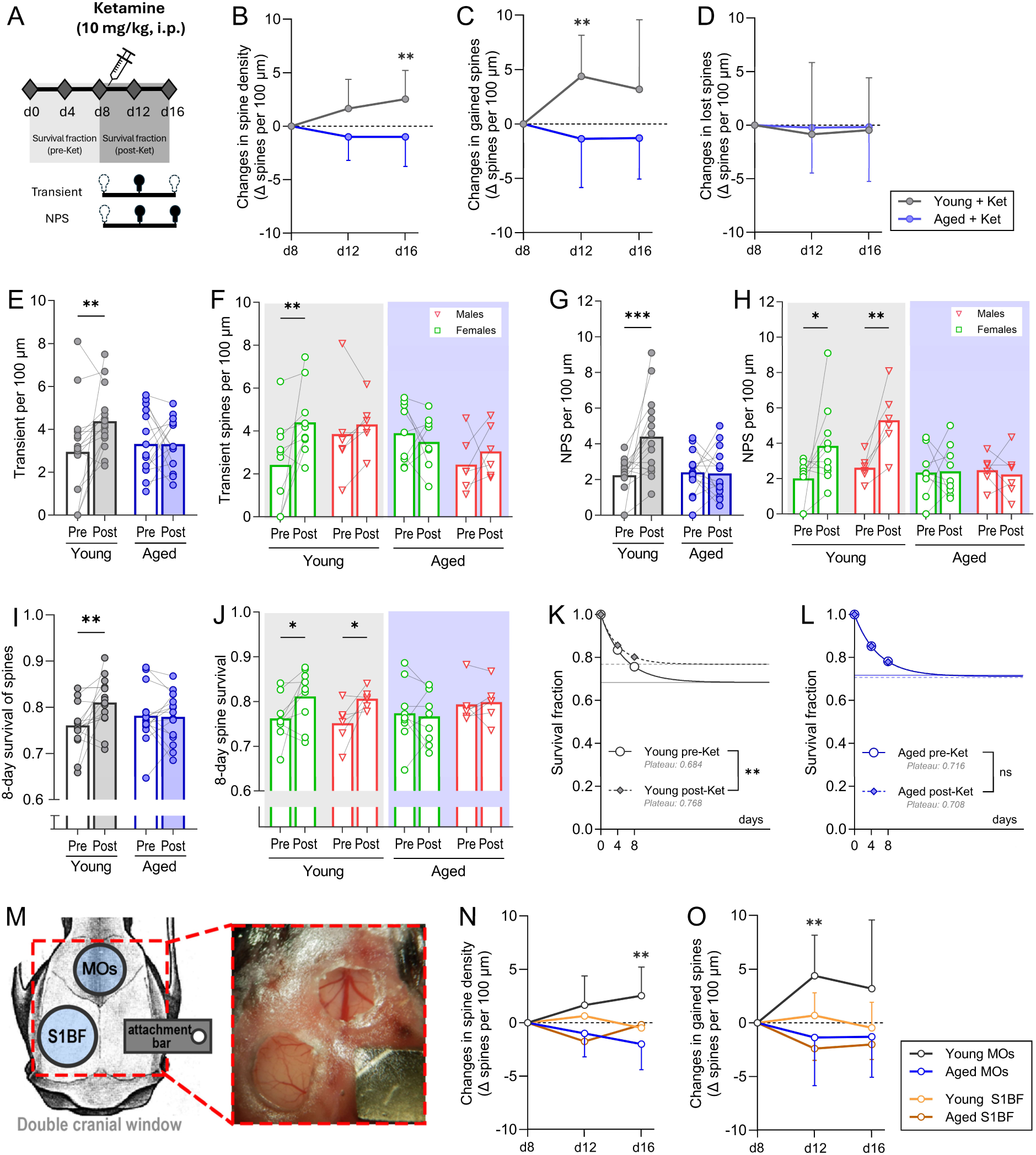
**A** Schematic of the in vivo 2PE imaging timepoints (diamonds). **B** Dendritic spine density of layer 5 pyramidal neurons in the MOs significantly increased at d16 compared to d8 in young but not in aged mice (Two-way ANOVA: Age x time F(1558, 45.19) = 5.648 *p* = 0.018; time: F(1558, 45.19) = 22.53, *p* < 0.0001; Tukeýs test young d16 vs d8: p = 0.048). **C** Gained spines is significantly incremented after the treatment with ketamine in young but not in aged mice (Two-way ANOVA: Age x time F(1750, 50.75) = 5.648 *p* = 0.0104; time: F(1750, 50.75) = 1.354, *p* = 0.0007; Tukeýs post-hoc young d12 vs d8: p = 0.0009). **D** Lost spines as an effect of ketamine treatment show no differences as a consequence of treatment or age. **E** Transient spines increase as a consequence of ketamine in young but not aged mice (Two-way ANOVA: age x treatment ns; Fisheŕs LSD young pre-Ket vs post-Ket: *p* = 0.007), **F** Transient spines increase specifically in young males compared to young females (Two-way ANOVA: sex x treatment F(1,14) = 3.484, *p* = 0.08; Fisheŕs LSD male pre-Ket vs post-Ket: *p* = 0.017), being this effect absent in aged mice. **G** New persistent spines increased in young compared to aged mice (Two-way ANOVA with repeated measurements: age x treatment F(1,29) = 8.736, *p* = 0.061; Tukeýs test young pre-Ket vs post-Ket: *p* = 0.0009). **H** New persistent spines in young male and female mice are equivalently increased by ketamine treatment (Two-way ANOVA: sex x treatment F(1,14) = 0.4418, *p* = 0.523; Fisheŕs LSD male pre-Ket vs post-Ket: *p* = 0.0345; female pre-Ket vs post-Ket: *p* = 0.019). **I** 8-day spine survival is improved after ketamine treatment in young but not aged mice (Two-way ANOVA with repeated measurements: age x treatment: ns; Fisheŕs LSD young pre-Ket vs post-Ket: *p* = 0.015). **J** 8-day spine survival separated by sexes shows a similar effect of ketamine-induced increased in survival in both males and female mice (Two-way ANOVA: sex x treatment F(1,14) = 0.0277, *p* = 0.87; Fisheŕs LSD male pre-Ket vs post-Ket: *p* = 0.0417; female pre-Ket vs post-Ket: *p* = 0.0452). **K** Estimated fraction of persistent spines in young mice is increased after ketamine administration (F 4.95(2, 78) *p* = 0.0094, plateau 0.68 pre-Ket vs 0.77 post-Ket; *K* = 0.1835 pre-Ket vs 0.24 post-Ket). **L** Estimated fraction of persistent spines is unaltered in aged mice after ketamine administration. **M** Representative image of the locations of the cranial windows. **N** Spine density and **O** gained spines after ketamine treatment do not change in the somatosensory cortex of young or aged mice. Density, dynamics, and survival fraction in MOs: n=9/6 neurons from 7/5 young mice, and n= 9/6 from 6/5 aged mice (male/female respectively, in both cases). Density and gained spines in S1BF: n= 13 neurons from 7 young mice and n= 8 neurons from 5 aged mice. Data presented as mean ± SD.

Given that ketamine did not affect spine density or dynamics in aged mice, we asked whether ketamine could enhance the stability of pre-existing spines. To test it we measured the 8-day survival fraction and estimated long-term spine survival before and after ketamine treatment. While ketamine significantly increased the 8-day spine survival as well as the predicted fraction of persistent spines (Fig. 4I-L), it had no such effect in aged mice, suggesting that the drug fails to enhance the persistence of existing spines in the frontal cortex during aging.

Moreover, to test whether ketamine elicits a global spinogenic effect in the brain, or whether the effect is region-dependent, we assessed ketamine-induced plasticity in a different brain region, the S1BF. For this, we implanted a double cranial window in some of the animals, placing one on the MOs and a second cranial window on the somatosensory cortex (Fig. 4M). We found that ketamine was unable to elicit plasticity in the somatosensory cortex of neither young nor aged mice, indicating that the effect of ketamine on structural synaptic plasticity is region-specific (Fig. 4N-O).

## Discussion

In this study we used a longitudinal *in vivo* imaging approach to investigate whether synaptic dynamics change with aging in the frontal cortex of mice. Changes on the density of dendritic spines have been reported to vary depending on the region and the cell type^39,40^ using histological preparations in fixed tissue. In this specific region, we found that steady-state synaptic density and dynamics in the apical shaft of the layer 5 pyramidal neurons are preserved through aging. These results are in agreement with previous studies performed by our team in other cortical regions, where we found similar trends towards the preservation^35^, or even an increase^18,34^, of the density and dynamics of dendritic spines of these neurons with aging.

An interesting finding of this study is that the observed sex differences in dendritic spine density and dynamics in the frontal cortex of young mice, with female mice displaying more dendritic spines, but more stable, than male mice, are preserved through aging. There is a growing body of evidence reporting sex differences in density at several life periods and in different parts of the brain^41–44^. However, the temporal dynamics and stability of dendritic spines in the frontal cortex have not been widely explored. Here we report that layer 5 pyramidal neurons of female mice have a significantly higher fraction of stable dendritic spines than males leading to significantly higher fractions of persistent spines in both young and aged mice. Interestingly, we did not find differences with aging in the fraction of stable vs unstable spines in female mice, indicating that those differences are not likely due to hormonal state and estrous cycle phase, since they are preserved in aged female mice.

We utilized ketamine, a non-competitive NMDA receptor antagonist, to induce structural synaptic plasticity, as it has been previously reported to induce spinogenesis in this region^29^. Ketamine has emerged as a promising treatment for patients with treatment-resistant depression, paving the way for the use of dissociative agents in the management of psychiatric disorders^45,46^. Although the present study was not designed to evaluate differences in the antidepressant efficacy of ketamine in the aged prefrontal cortex, our findings provide some insights into how age-related factors may modulate its therapeutic effectiveness. In this regard, the SUSTAIN-2 clinical trial proved the effectiveness of intranasal esketamine combined with oral antidepressants in adult patients^45^. However, when the same dosing regimen was applied in the TRANSFORM-3 trial targeting elderly patients with treatment-resistant depression, the antidepressant effect was not replicated in the same way^47^. A post-hoc analysis revealed that patients in the range of 65-74 years old of the TRANSFORM-3 study exhibited partial improvement, whereas patients over 75 years old failed to respond to treatment to the same extent^46^. Supporting this decreased of ketamine’s efficacy with aging, another recent study demonstrated that the antidepressant effect of ketamine was short-lived in older adults^48^, which contrasts with the longer effect described in young individuals^45,46^. Taken together, our findings suggest that the MOs of aged mice exhibits a diminished capacity to express structural synaptic plasticity compared to those of young mice, synaptic plasticity deficit that may underlie the reduced efficacy of plasticity-based antidepressant treatments such as ketamine. Further studies are needed to elucidate the mechanisms driving this age-related synaptic plasticity impairment and to identify strategies that can enhance therapeutic responses in aging populations. Interestingly, we did not find overall sex-differences in the spinogenic effect of ketamine, indicating that its effectiveness may not be sex-dependent.

It is interesting to note, however, the differences observed between brain regions in ketamine’s ability to induce structural plasticity of dendritic spines in young mice. In this study, while MOs shows an increase in dendritic spine density and survival after ketamine treatment, S1BF is unaltered. Our interpretation is that ketamine acts through several synergistic mechanisms to induce spine plasticity, and only when those mechanisms converge is when the spinogenic effect is achieved. In this regard, it has been described that ketamine increases excitability of layer 5 pyramidal neurons through disinhibition, which increases glutamate release^28^. At the same time, ketamine’s ability to induce formation of dendritic spines has been demonstrated to be dependent on dopaminergic mechanisms^49^ and that dopamine release is necessary to induce formation of new dendritic spines. The co-release of glutamate and dopamine has been described in the frontal cortex^50,51^ a brain area that receives a higher number of dopaminergic projections, mostly from the ventral tegmental area, than the somatosensory cortex^52,53^. Therefore, we believe that these region-specific innervation differences are responsible for the ability of ketamine to trigger structural synaptic remodeling, mainly manifested as an increase in the formation of dendritic spines first, followed by a higher stabilization rate of these newly formed spines.

The deficits in ketamine-induced remodeling of dendritic spines of layer 5 pyramidal neurons may be an indirect indication of the underlying impairments in synaptic plasticity occurring in the aged prefrontal cortex and that may also be responsible for the impairments in cognitive flexibility observed in the four-odor choice discrimination and reversal task in the aged group. Our behavior data are in agreement with other studies showing decreased behavior flexibility in mice^54–56^ and humans^57–60^ as a consequence of aging. Despite substantial evidence that aging impairs cognitive flexibility, few studies have investigated the underlying mechanisms, and even fewer have addressed how aging affects synaptic dynamics in frontal cortical regions. In this context, our selection of the four-odor choice discrimination and reversal task is particularly relevant, as it directly engages the MOs, a brain region in which synaptic plasticity has been shown to be a critical determinant of performance in this task^32^, which was the main reason to drive this study in the effect of aging in the MOs region.

Our results from longitudinal imaging of dendritic spines are limited to layer 5 pyramidal neurons, yet there may be differences regarding the impact of aging on different cortical layers and cell types. Relevant to this study, layer 5 pyramidal neurons can be further classified based on their projection targets, distinguishing intratelencephalic (IT) neurons, which project within the cortex and striatum, and pyramidal tract (PT) neurons, those that project to subcortical structures. These layer 5 neuron subtypes not only serve distinct functional roles but also exhibit differential dendritic spine densities^61^ as well as differential vulnerabilities to neurological diseases^62^. The GFP-positive layer 5 pyramidal neurons of Thy1-GFP-M mice imaged in the present study are projection neurons^30^, therefore classified within PT neurons, as we have also previously reported using their intrinsic and synaptic electrophysiological properties^63^. There is no available data on density and dynamics of dendritic spines of layer 5 IT pyramidal neurons This raises the possibility that PT and IT neurons may exhibit differential vulnerabilities to aging, which could contribute to the inconsistencies reported in the literature regarding the effects of aging on synaptic plasticity in layer 5 pyramidal neurons: while some studies have described increases in basal spine density and dynamics with aging^18,34,64^, others have reported decreased densities^15,16,65^, or no significant changes^27^. Furthermore, it is important to acknowledge that spine density and dynamics are strongly influenced by species, brain region, sex, neuronal compartment, and the methodological approaches used for staining, tissue processing, and data collection and quantification. Collectively, these factors contribute to substantial inter-study variability and prevent the creation of broadly generalizable principles in aging studies.

Elucidating changes in dendritic spine plasticity of deeper frontal cortical regions, such as prelimbic or infralimbic prefrontal cortex, would broaden our understanding of the degree of plasticity that these higher order brain areas of the medial prefrontal cortex are able to express. However, there are technical limitations when attempting to reach medial and deeper prefrontal cortical areas using the approach of cranial window implantation and 2PE imaging. As the current state-of-art, imaging at the single dendritic spines resolution in prelimbic or infralimbic prefrontal cortex can only be achieved by the use of three-photon excitation microscopy^66,67^. The use of implanted microprisms has been successful in mice to reach intermediate regions such as cg1, but not prelimbic or infralimbic regions of the medial prefrontal cortex^68^.

## Conclusion

Altogether, our data reveal that the synaptic architecture of layer 5 pyramidal neurons is mostly preserved through aging in the MOs region of healthy aging mice, contrasting with the idea of significant synaptic loss that is thought to be a firm statement in the field of aging^69,70^. Despite the preservation of dendritic spines in the prefrontal cortex, the ability of the aged circuits to remodel connections in response to external stimuli may be compromised, leading to impairments in behavior flexibility. This may explain why interventions that rely on boosting synaptic plasticity, such as the use of ketamine for treatment of treatment-resistant depression, fail to achieve full efficacy in older individuals. Developing strategies that restore or bypass these plasticity mechanisms through co-administration of plasticity-priming agents, targeted circuit interventions, or age-specific dosing protocols, may be necessary to extend the therapeutic benefits of pharmacological agents across lifespan.

## Acknowledgements

This work is supported by National Institutes of Health (NIH) NINDS R R01NS114286, and NIA R01AG074489 to RM.

---

**Supplementary figure 1.**
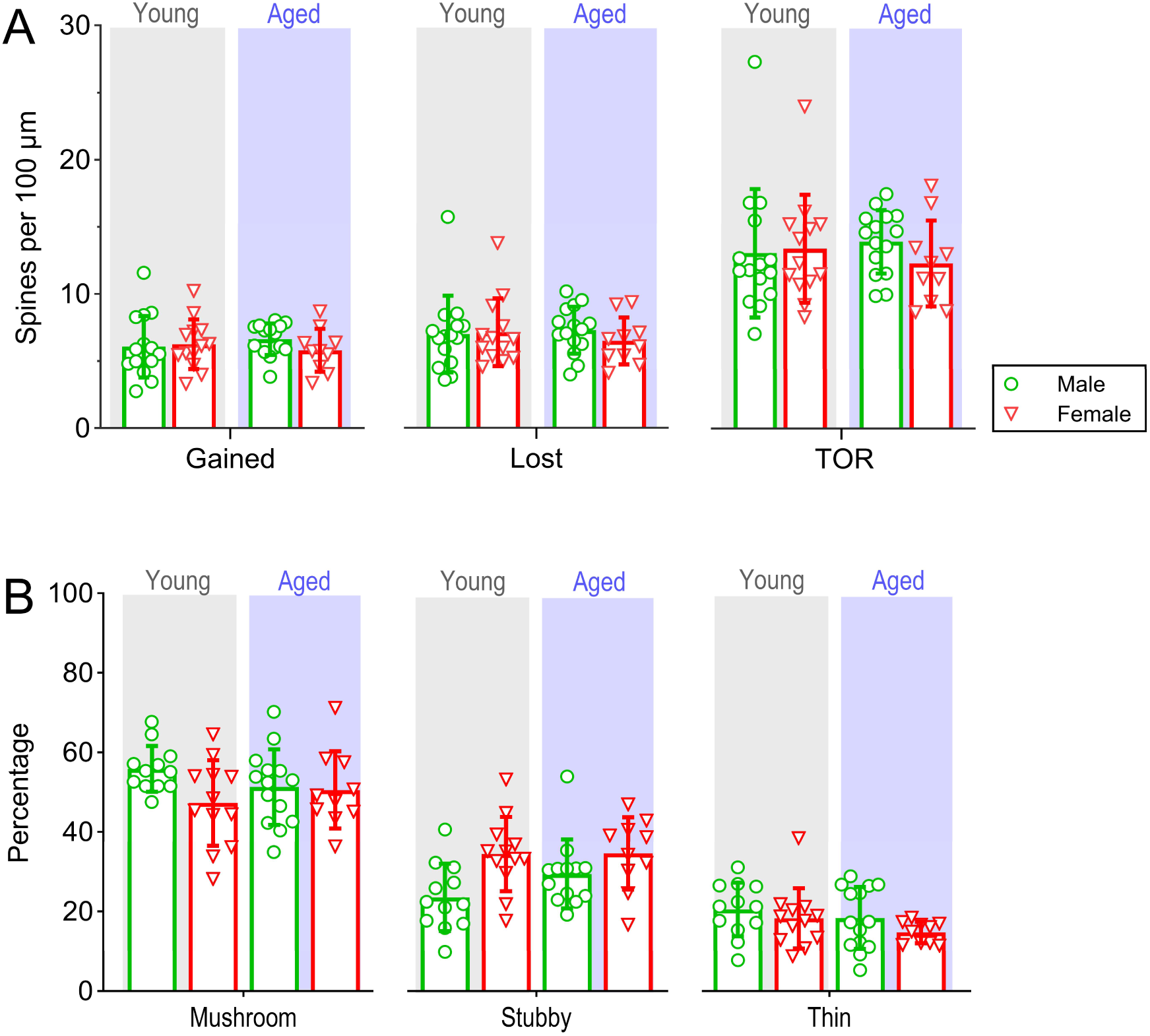
**A** Steady-state dendritic spine density and dynamics represented in absolute values (spines per 100 μm) in young and aged males and females. **B** Percentage of the different type of morphologies of dendritic spines (mushroom, stubby, thin) per neuron. n = 31 neurons from 19 young mice, and n = 25 neurons from 15 aged mice. Data presented as mean ± SD.

**Supplementary figure 2.**
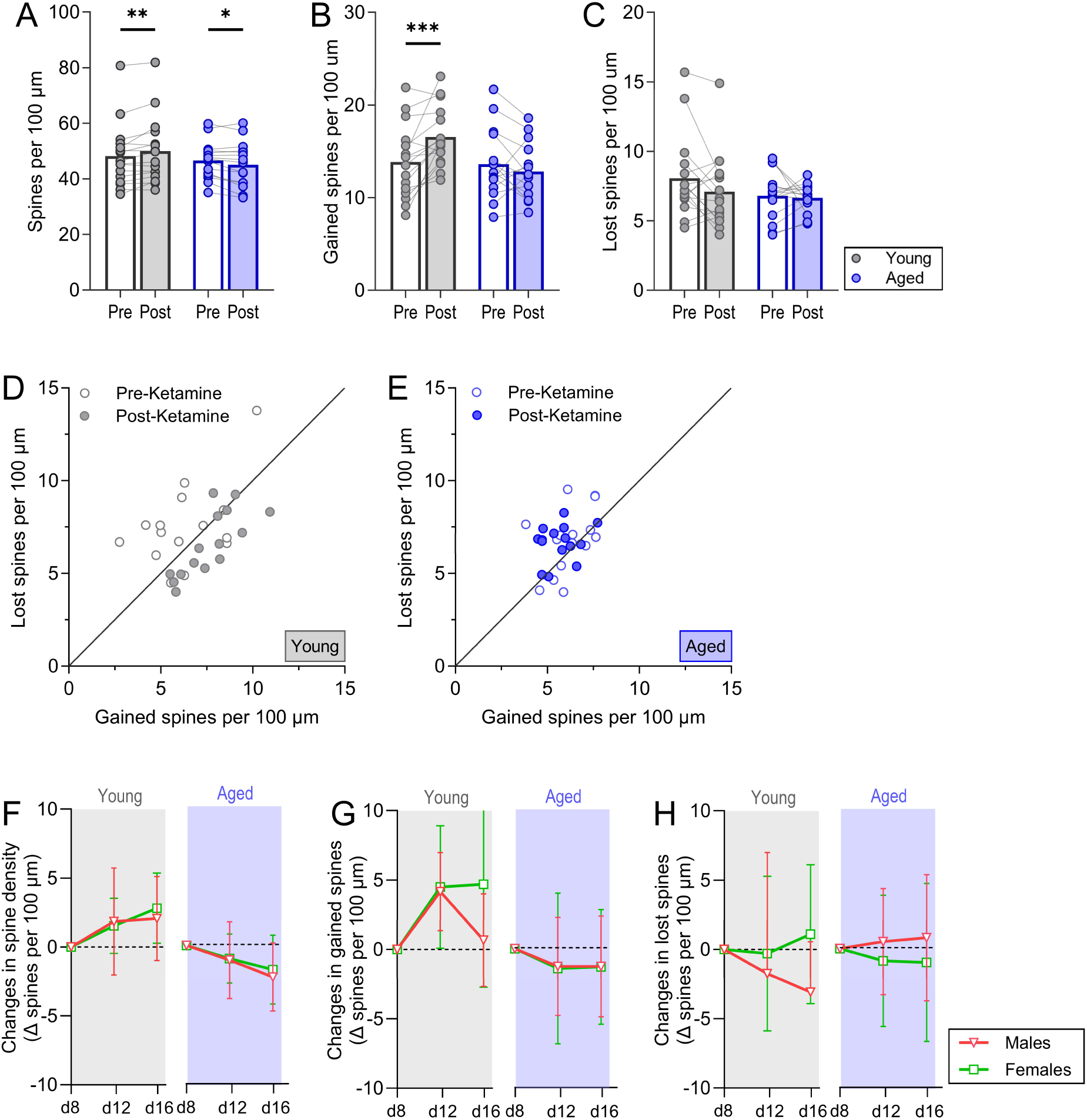
**A** Dendritic spine density, **B** gained spines, **C** and lost spines pre- and post-ketamine administration (spines per 100 μm) in young and aged mice. Representation of the balance gained/lost per neuron in **D** young and **E** aged mice. Sex differences in ketamine-induced changes in **F** density, **G** gained spines, **H** and lost spines with respect to day 8.

